# Redox fluctuations control the coupled cycling of iron and carbon in tropical forest soils

**DOI:** 10.1101/312108

**Authors:** Amrita Bhattacharyya, Ashley N. Campbell, Malak M. Tfaily, Yang Lin, Whendee L. Silver, Peter S. Nico, Jennifer Pett-Ridge

## Abstract

Oscillating redox conditions are the norm in tropical soils; driven by an ample supply of reductants, high moisture, microbial oxygen consumption, and finely textured clays that limit diffusion. Yet the net result of variable soil redox regimes on iron-organic matter (Fe-OM) associations in tropical soils owing to changing climate is poorly understood. Using a 44-day redox incubation experiment with humid tropical soils from Puerto Rico, we examined patterns of Fe and C transformation under four redox regimes: static anoxic, flux 4-day (4d oxic, 4d anoxic), flux 8-day (8d oxic, 4d anoxic) and static anoxic. Prolonged anoxia promoted reductive dissolution of Fe-oxides and an increase in short-range ordered (SRO) Fe oxides. Preferential dissolution of this less-crystalline Fe pool was evident immediately following a shift in bulk redox status (oxic to anoxic), and coincided with increased dissolved organic carbon, presumably due to acidification or direct release of OM from dissolving Fe(III) mineral phases. Average nominal oxidation state of water-soluble carbon was lowest under persistent anoxic conditions, suggesting more reduced OC is microbially preserved under reducing conditions. Anoxic soil compounds had high H/C values (similar to lignin-like metabolites) whereas oxic soil compounds had higher O/C values, akin to tannin- and cellulose-like components. Cumulative respiration derived from native soil organic carbon was highest in static oxic soils. These results highlight the volatility of mineral-OM interactions in tropical soils, and suggest that short-term impacts of shifting soil O2 availability control exchanges of C between mineral-sorbed and aqueous pools, implying that the periodicity of low-redox moments may control the fate of C in wet tropical soils.

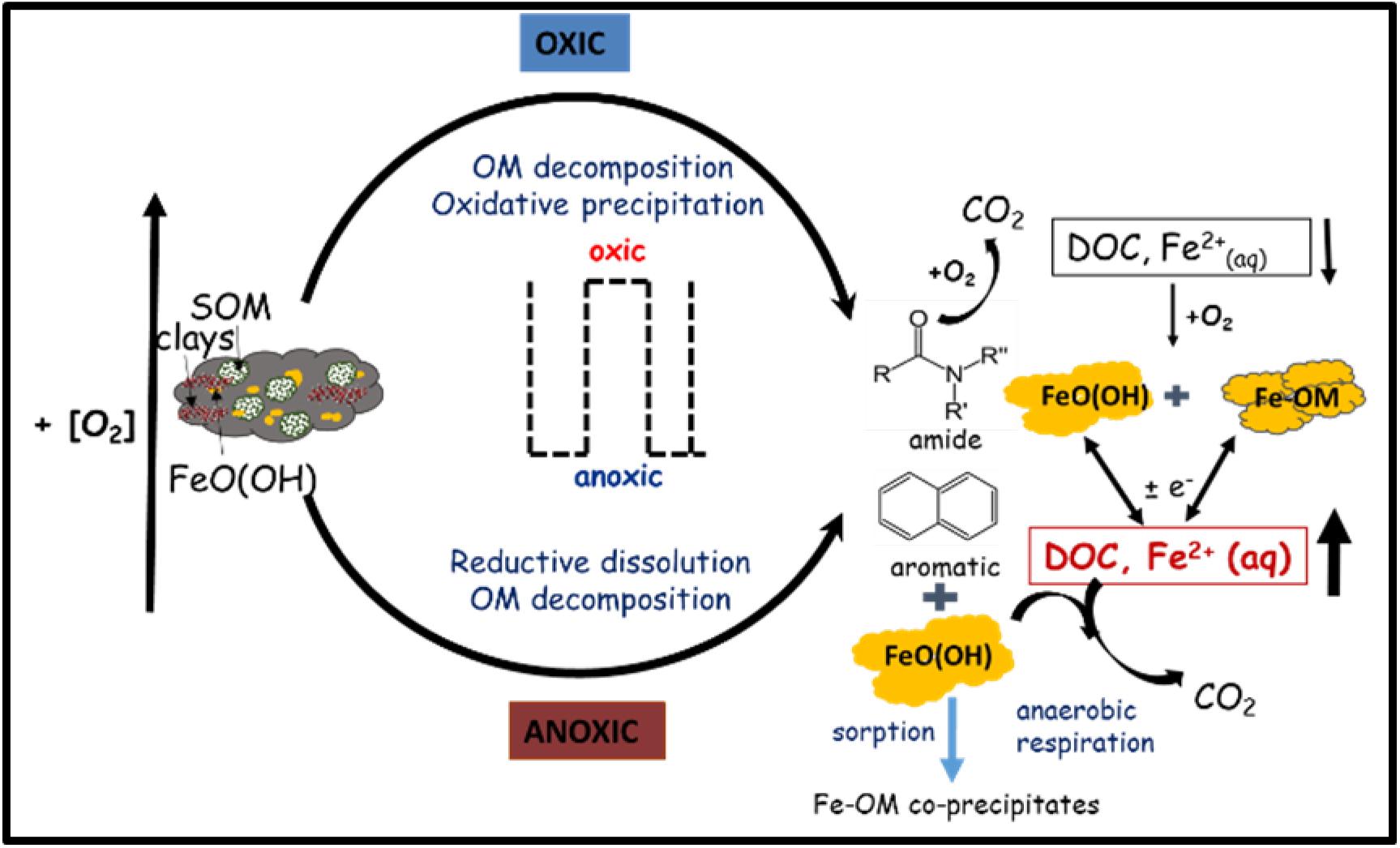
Toc Art.

## INTRODUCTION

Wet tropical soils frequently alternate between fully oxygenated and anaerobic conditions, making them biogeochemical ‘hotspots’ for redox reactions.^1-6^ In the wet tropics, soils are often dominated by iron (Fe) (and aluminum) rich clays and oxides that sorb organic carbon (C), but are also susceptible to redox-induced mineral transformations.^6-10^ The rapid Fe and C cycling typical of wet tropical ecosystems is fueled by a characteristically dynamic redox environment driven by high biological oxygen demand, high moisture (limiting O2 diffusion), warm temperatures and abundant labile C. Unlike classically defined low-redox soils (e.g. wetlands)^11^ where redox zonation can be relatively static in time and space, redox oscillations in upland tropical forest soils are spatially and temporally heterogeneous, likely occurring at the individual aggregate scale and responding rapidly (hourly to daily)^6, 12, 13^ to variations in rainfall and dissolved organic carbon (DOC) inputs.

Iron minerals play a critical role in the C cycle of highly weathered tropical soils, particularly where frequent cycles of reduction and oxidation fuel a large portion of decomposition (~50% of C oxidation)^1-3, 14-16^ via a large Fe reducer population (up to 10^9^ per gram) that may be responsible for ~ 40% of soil CO_2_ respiration.^10^ Iron minerals also mediate belowground C retention ^4, 17-21^ by binding OM in surface complexes and aggregates.^8, 22-24^ This is particularly true for high surface area, short-range-ordered (SRO) Fe minerals, which are associated with some of the oldest C in terrestrial systems.^25, 26^ Much of the soil organic carbon (SOC) which is stabilized via Fe-OM associations is susceptible to redox effects because Fe can easily solubilize and reprecipitate/crystallize in response to local Eh conditions.^6^ In an oscillating redox environment, when O2 is often depleted, Fe minerals quickly become a favorite terminal electron acceptor and are reduced. This microbially mediated process can then liberate associated material (nutrients or OM) ^27, 28^, and we expect this liberated OM to rapidly appear in the DOM pool. Previous work in Hawaiian tropical soils suggests Fe mineralogy evolves more quickly to a crystalline endpoint when soil redox is variable, due to preferential dissolution of less crystalline materials during repeated reduction phases.^6, 8^ High-surface area SRO Fe phases are likely produced by rapid mineral precipitation events and localized at redox interfaces in the soil profile. Yet, these same mineral phases are susceptible to rapid Fe reduction when O2 becomes limiting, and this in turn may lead to C oxidation and increased C mobility (via colloid dispersion).^7, 29^

In the coming 50 years, our study site, the US Forest Service Luquillo Experimental Forest (LEF) (within the El Yunque National Forest of northeastern Puerto Rico), is predicted to experience shifts in rainfall and temperature^30^ that may significantly affect both soil moisture and the periodicity of soil redox oscillations. Multiple microbial/mineral driven processes could be affected, including OM degradation, methanogenesis, denitrification, and Fe reduction/oxidation^31-39^, since soil moisture regulates these key redox-sensitive aspects of the soil C cycle.^4^ We expect that a shift to warmer, drier, more oxygenated soil conditions would strongly impact mineral-organic interactions (e.g. Fe-OM associations).^6-9, 35, 36, 40^ However, the degree to which Fe-oxides associate with SOM in tropical forest soils, particularly under fluctuating redox conditions is poorly understood, and little is known about how adsorbed and co-precipitated SOM impact biological Fe reduction and secondary Fe mineralization. What may be far more important than mean Eh (or mean soil [O2]), is the *pattern* of redox oscillation, *e.g.* its periodicity (the recurrence rate of low-redox events) and duration (the persistence of low or high redox periods).

A better understanding of the influence of redox variability on Fe-OM relationships would aid in a better designing of the tropical C cycle. Hence, in this study we sought to understand the impact of fluctuating redox conditions on Fe-OM interactions on wet tropical soils which were subjected to episodic oxidizing/reducing events using a replicated, high temporal resolution redox oscillation study. We test the hypothesis that the periodicity and duration of redox regimes will alter Fe speciation together with the chemical composition of water extractable OM. Our results show how the nature of Fe-OM interactions change in response to fast redox switches which, in turn, impact the coupled Fe-C biogeochemical cycle in tropical terrestrial soils.

## MATERIALS AND METHODS

### Study site and soil sampling

Our study site in the LEF is part of a Critical Zone Observatory and Long-Term Ecological Research site (Lat. 18° 18 N; Long. 65°50’ W). The Luquillo Mountains have steep topography, with distinct gradients in rainfall and temperature, and naturally dynamic soil redox regimes.^5^ Highly weathered oxisols occur on parent material derived from volcaniclastic minerals (clay rich) that experience frequent soil O2 depletion (mean O2 is 13 ± 0.21 %, oscillating between 3 and 17 %)^4, 5^ with major shifts every 4 – 6 days.^4, 5^ The mean water content is 35-65 % and pH is ~ 4.8.^41^ Notably, LEF soils are enriched in SRO mineral phases that are highly susceptible to Fe reduction. Surface soils (0-10 cm) for our study were collected from a slope location near the El Verde field station in LEF in late January 2016 and shipped overnight to Lawrence Livermore National Laboratory, CA at ambient field temperature and moisture. Once in the laboratory, soils were gently homogenized and visible plant debris, rocks, and soil macro-fauna were removed manually.

### Experimental Design

Our experimental scheme for manipulating bulk soil redox followed previously established methods ^40^; approximately 20 g (oven dry weight) of soil was weighed into a 500 mL glass microcosm, and a total of 128 replicate microcosms (to our knowledge, the largest, most highly replicated experiment of its kind) were established. Microcosms were subjected to a 16-day pretreatment period to overcome the disturbance effects that might have occurred due to soil handling and homogenizing. Microcosms were capped with gas-tight lids fitted with a GeoMicrobial septa (Geo Microbial Technologies, Ochelata OK) with tygon tubing extending into the interior of the jar. Oxygen (oxic) or nitrogen (anoxic) gases were delivered via the tubing to the bottom of each jar at a rate of 3 ml/min and vented with a syringe needle in the septa. All microcosms were initially treated with 4 days of oxic conditions by flushing the headspace with ambient air, followed by 4 days of anoxic flushing with N2 (one complete redox cycle),and repeated this cycle one more time before the first harvest. Microcosms were then divided into four redox regime treatment categories: (1) static anoxic, (2) static oxic, (3) 4 days oxic/4 days anoxic (flux 4-day), (4) 8 days oxic/4 days anoxic (flux 8-day) and incubated for 44 days, with 3 replicates per treatment and harvest point (5 replicates for the final harvest). Headspace was manipulated in the same way as in the pre-treatment. Flux 4-day and flux 8-day treatments started and ended under an oxic incubation headspace.

### Microcosm Harvesting

During the 44-day experiment, replicate microcosms from all four redox treatments were destructively harvested on days 12, 20, 23, 36 and 44. Additionally, flux 4-day and flux 8-day treatments were harvested on days 16 and 33 respectively. For flux 4-day and flux 8-day treatments, microcosms were harvested with a more intensive time schedule--immediately prior to the headspace gas switch (0 minute), at 30 minutes and 180 minutes--in order to capture shortterm geochemical responses as a function of headspace gas composition. During the harvest, soils were lightly homogenized with a spatula after opening the microcosm jar and then subsampled for microbial and chemical analyses. Sampling and extractions for microcosms assigned to an anoxic headspace were performed within an anoxic glove box chamber (5% H2 and 95% N2). All the chemical reagents used within this anaerobic chamber were prepared with degassed water to preserve the oxidation state of Fe.

### Geochemical Methods

#### Amorphous Fe by Acid Ammonium Oxalate Extraction

Soils were treated with a selective dissolution procedure to estimate the amount of Fe in operationally defined “amorphous” or “short range ordered” fractions, and extracted with ammonium oxalate in the dark to assess for Fe in poorly-crystalline or “amorphous” inorganic materials. For the acid ammonium oxalate (AO, Feo) extraction^42^, 0.25 g soil (dry weight) was weighed into foil-covered centrifuge tubes, 10 ml of the oxalate solution (0.2 M of acid ammonium oxalate solution at pH ~ 3) was added, and the suspensions were shaken for 4 hours. The suspensions were then centrifuged at 10,000 rpm for 20 min, and the supernatants filtered (0.45 μm). Extracts were analyzed for Fe by Inductively Coupled Plasma Atomic Emission Spectroscopy (ICP-AES) at Lawrence Berkeley National Laboratory.

#### Determination of Ferrous Iron (Fe^2+^) by 0.5M HCl Extraction Method

To measure ferrous iron (Fe^2+^), 1.0 g soil (dry weight equivalent) was weighed inside the anaerobic chamber and an aliquot of 5mL of 0.5M HCl was added to digest the samples. The mixture was swirled for 30s and left in the dark overnight to ensure complete digestion.^43^ The supernatant was then filtered using a 20μm cellulose acetate filter. Fe(II) in the extract was measured colorimetrically using the ferrozine method^43^; 100 uL or 10 uL (when [Fe(II)] > 1 mmol L^−1^) of the filtrate was added to 0.5 mL ferrozine solution (1 g/L ferrozine in 50 mmol/L HEPES buffer, pH 7.0) along with 4 ml of MilliQ (18.2Ω) water. After 10 minutes, absorbance was measured at a wavelength of 562 nm with a UV-Vis spectrophotometer (Thermo Scientific). The concentration of ferrous (Fe^2+^) iron was calculated and reported in g Fe^2+^ (kg soil)^−1^.

#### Fe speciation by X-ray absorption spectroscopy

Iron speciation (oxidation state and chemical coordination environment) was determined using X-ray absorption spectroscopy (XAS) analyses. Fe K-edge extended X-ray absorption fine structure (EXAFS) (7112 eV) was conducted on beamline 4-3 at the Stanford Synchrotron Radiation Lightsource (SSRL), at Menlo Park, CA, under ring operating conditions of 3 GeV with a current of 450 mA. Due to the limited time available at the beamline only samples from the pretreatment and final harvest days were chosen to document the change in Fe speciation due to redox incubation. Samples were sealed on Teflon holders with Kapton tape inside an anaerobic glove bag to prevent oxidation during sample preparation. Samples were then mounted in a N2 (l) cryostat to preserve the oxidation state of Fe and to prevent potential beam damage which might occur during data collection. A double crystal Si (220) monochromator with an unfocused beam was detuned 30% to reject harmonics affecting the primary beam. Between 7 and 10 individual spectra were averaged for each sample. Pure elemental Fe foil was used for Fe energy calibration at 7112 eV. A Lytle detector was used to record the fluorescence spectra of EXAFS and XANES scans. Scans were deadtime corrected, and averaged using SixPack software.^44^ The fluorescence spectra were averaged and pre- and post-edge subtracted using the Athena software package.^45^ Background removal, normalization, and glitch removal were also performed in Athena. Linear combination fitting (LCF) of spectra was performed in Athena^45^ in k^3^-weighted k-space between k = 2 and 12, using the following end-members: siderite (FeCO3), 2-and 6-line ferrihydrite [Fe(OH)_3_.nH_2_O], goethite (α-FeOOH), lepidocrocite (γ-FeOOH), hematite (α-Fe2O3), pyrite (FeS2), mackinawite (FeS), magnetite (Fe3O4) and illite. These references were chosen based on their likelihood to be present in our experimental samples. Compounds were only included in the fit if the factional contribution greater than 0.05. Additional details for the EXAFS analysis are presented in the Supporting Information.

#### Estimation of water extractable dissolved organic carbon (DOC) content

Water extractable DOC was extracted by shaking 2.0 g of soil (dry weight equivalent) in 200 ml Milli-Q water (18.2 Ω) for 1 h at 80 rpm. Suspensions were then filtered through 0.45 μm Nylon membrane filters using a vacuum filtration technique. The filtrate was then analyzed for dissolved organic carbon using a Shimadzu (Japan), Total Organic Carbon Analyzer (TOC-vcsh) at Lawrence Berkeley National Laboratory.

### FT-ICR-MS data acquisition and data analysis

#### Soil Sample Preparation

Samples were prepared according to Tfaily et al. 2017.^46^ Briefly, 1 mL of 18.2 MΩ Milli-Q water (EMD Millipore, Billerica, MA) was added to 300 mg bulk soil, vortexed for 20 sec and then put onto an Eppendorf Thermomixer at 1000 rpm and 20°C for 4 hours. Samples were then removed from the mixer and spun down before the supernatant was pulled off frozen and the final 1 mL chloroform was added and the samples were shaken overnight was described above. This process was repeated for all the samples.

#### Organic carbon characterization

Ultrahigh resolution characterization of water extractable DOC was carried out using a 12 Tesla Fourier-transform ion-cyclotron-resonance mass spectrometer (FT-ICR-MS) (Bruker SolariX, Billerica, MA). Water extracts were thawed overnight at 4°C and then diluted with methanol to improve electrospray ionization efficiency, no precipitation was observed. Suwannee River Fulvic Acid (SRFA) standard, obtained from the International Humic Substance Society (IHSS), was used as control and injected after every 20 samples followed by a methanol blank to assure instrument stability and to verify no sample carryover. Samples were injected directly into the instrument using an automated custom built system, and ion accumulation time was optimized for all the samples, depending on extraction solvent. Optimal parameters were established prior to the samples being injected.^47-49^ Samples were introduced to the electrospray ionization ESI source equipped with a fused silica tube (30 um internal diameter) through an Agilent 1200 series pump (Agilent Technologies), at a flow rate of 3.0 uL min^−1^. Experimental conditions were as follows: needle voltage, +4.4 kV to generate negatively charged molecular ions; Q1 set to 50 m/z; 144 individual scans were co-added for each samples and internally calibrated using OM homologous series separated by 14 Da (–CH2 groups). The mass measurement accuracy was typically within 1 ppm for singly charged ions across a broad m/z range (100-1100 m/z). To further reduce cumulative errors, all sample peak lists for the entire dataset were aligned to each other prior to formula assignment to eliminate possible mass shifts that would impact formula assignment. Putative chemical formulas were assigned using Formularity software.^50^ Chemical formulae were assigned based on the following criteria: S/N >7, and mass measurement error <1 ppm, taking into consideration the presence of C, H, O, N, S and P and excluding other elements. Peaks with large mass ratios (m/z values > 500 Da) often have multiple possible candidate formulas. These peaks were assigned formulas through propagation of CH2, O, and H2 homologous series. Additionally, to ensure consistent choice of molecular formula when multiple formula candidates are found the following rules were implemented: we consistently pick the formula with the lowest error with the lowest number of heteroatoms and the assignment of one phosphorus atom requires the presence of at least four oxygen atoms.

Nominal oxidation state of carbon (NOSC) was calculated using the stoichiometry of each assigned formula of each organic compound identified in the water extracts.^51^ The relevance of thermodynamic limitations under variable redox conditions can be evaluated by monitoring NOSC values as a function of the redox gradient. Furthermore, partial least square - discriminant analysis (PLS-DA) plots, generated by Metaboanalyst^52^, was used as a direct and rapid tool to optimize the separation of a particular class of metabolites based on the headspace composition on flux 8-day samples. Organic compounds that were significantly differnet between two treatments (i.e., organic compounds with a VIP (variable importance in projection) score >1 were further examined in van Krevelen diagrams and assigned to major biochemical classes based on the molar H:C (y-axis) and O:C (x-axis) ratios.^46^ These compounds were clustered into two groups (1) high O/C and H/C that it is on the carbohydrate region of the van-Krevelen diagram; and (2) compounds with medium O/C and low H/C that fit in the aromatic region of the van-Krevelen diagram.

##### CO_2_ flux measurements

Microcosm headspace gas samples for carbon dioxide (CO_2_) measurement were collected throughout the experiment. Samples were collected from the set of replicate microcosms which were to be harvested on day 44 (n = 5) with gastight syringes and stored in pre-evacuated 20-mL serum bottles sealed with butyl rubber septa (Geo-Microbial Technologies, Ochelata, Oklahoma, USA). Gas samples were collected immediately prior to switching microcosm headspace gas (redox state). CO_2_ concentrations were measured with a gas chromatograph (GC-14A, Shimadzu, Columbia, MD) equipped with a thermal conductivity detector.

## RESULTS AND DISCUSSION

### Long-term redox shifts--effects on Fe and DOC chemistry

Results from 0.5M HCl extractable Fe(II), oxalate extractable AO-Fe and water extractable DOC are presented in Figure 1. Soluble Fe(II) concentrations, which represent adsorbed and solid phase Fe(II)^53^, range from 0.06 – 2.17 g kg^−1^ soil (Figure 1a). Prolonged anoxia (static anoxic treatment) led to a significant accumulation of Fe(II) relative to other redox treatments (Figure 1a), whereas soils from fluctuating treatments had generally low Fe(II) concentrations and grouped closely with the static oxic Fe(II) values. Interestingly, the headspace gas composition had an impact on the Fe(II) content during the initial 23 days of the experiment for both the flux 4-day and flux 8-day treatments--soluble Fe(II) concentrations were higher when sampled under an anoxic headspace during the fluctuating treatments (days 16 and 23 for the flux 4-day, and days 12, 23 and 36 for the flux 8-day). Soils from both fluctuating treatments (days 12, 20, 36, 40 and 44 for flux 4-day; and days 20, 33 and 44 for flux 8-day) had similar Fe(II) concentrations to the static oxic soils when they were harvested during an oxic headspace exposure period. As expected, Fe(II) concentrations under static oxic conditions were consistently low, and ranged between 0.06 – 0.13 g kg^−1^ soil. Ammonium oxalate-extractable Fe (AO-Fe) (representing the amorphous or SRO Fe(oxy)hydroxide pool) was more than two times higher in static anoxic soils relative to the other treatments (Figure 1b), ranging from 3.8 – 10.2 g kg^−1^ soil. In soils incubated with fluctuating redox conditions, samples collected from an anoxic headspace period had higher amorphous Fe (5.6 g kg^−1^ at day 16) relative to those collected during an oxic period (4.5 and 4.8 g kg^−1^ at days12 and 20 respectively). Overall, AO-Fe concentrations in the fluctuating treatment soils grouped more closely with concentrations’ measured in the static oxic treatment soils. Water extractable DOC represents the organic C available for microbial respiration and in our incubations, ranged from 0.17 – 2.5 g kg^−1^ (Figure 1c). Concentrations were consistently high in the anoxic condition soils and were positively correlated with HCl-Fe(II) and AO-Fe concentrations. Soils collected during anoxic sampling points from the flux 4d and flux 8d treatments (days 16, 23 for flux 4d; days 12, 23 and 36 for flux 8d) had higher WEDOC concentrations relative to concentrations in oxic soils from the sampling points. However, towards the end of the experiment, the water extractable DOC concentrations for all the treatments converged around 0.44 g kg^−1^ soil—implying an overall decline in labile carbon availability.

**Figure 1.**
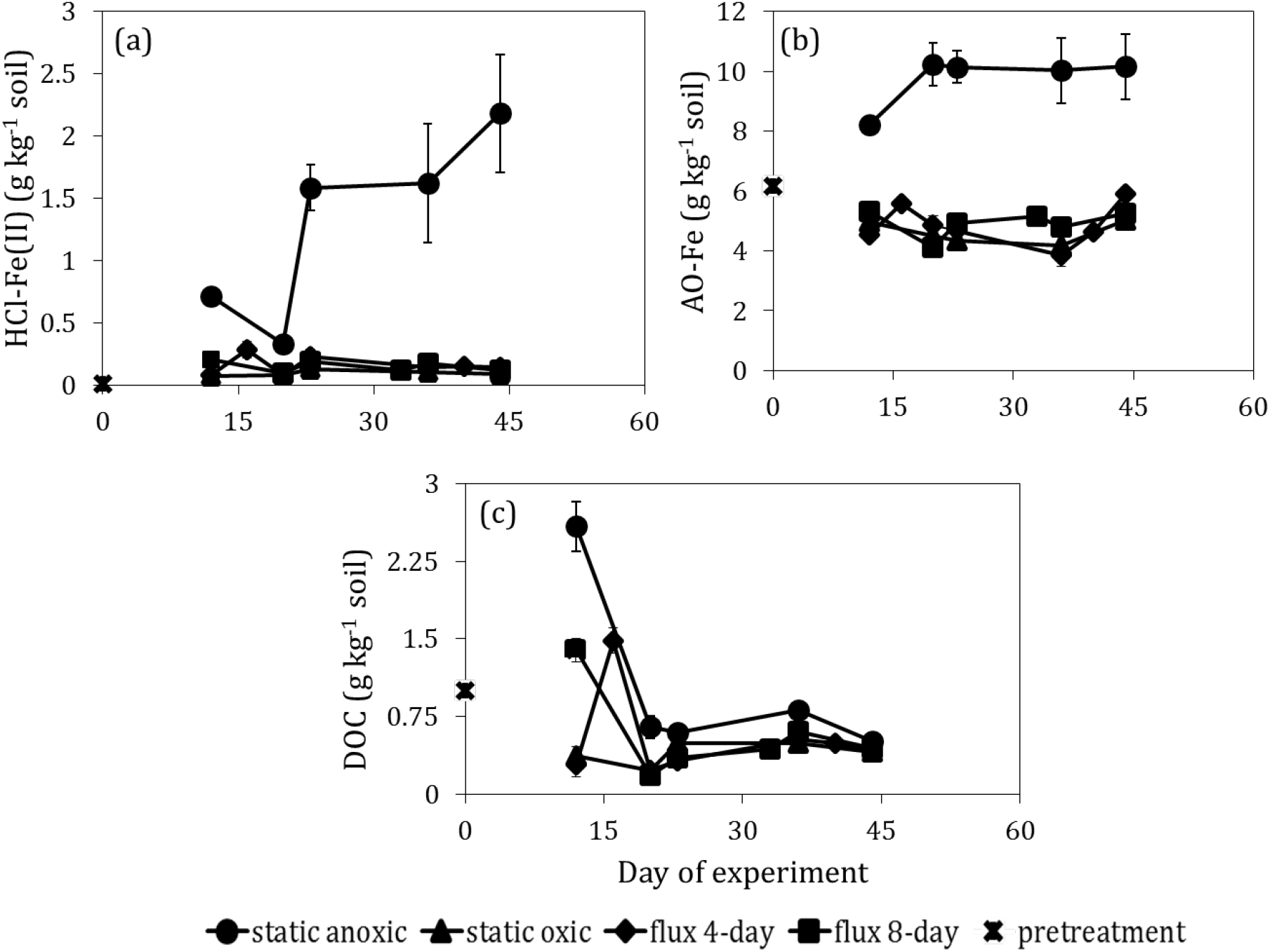
Changes in (a) 0.5N HCl extractable Fe(II); (b) oxalate extractable Fe (AO-Fe); and (c) water extractable dissolved organic carbon (WEDOC) concentrations (in g kg^−1^ soil) under four redox treatments over the 44-day incubation on a tropical forest soil from Puerto Rico. Concentrations for static anoxic, static oxic, flux 4-day, flux 8-day treatments are shown in blue, green, red and yellow respectively compared to that of the initial pretreatment concentration (black circle). Error bars indicate standard error for three replicate measurements.

In addition to the solution chemistry, linear combination fitting of the bulk Fe K-edge EXAFS spectroscopy provides semi-quantitative information on different Fe forms that could potentially be present under the different redox conditions (Table 1, SI Figure S1). Since soil samples from each redox treatment did not show any change in bulk Fe mineralogy over time, the data presented in Table 1 and SI Figure S1 are for samples harvested during pretreatment and at the end of the experiment (day 44). Irrespective of the redox condition, SRO Fe(oxy)hydroxide (ferrihydrite) was the dominant Fe species present in the soils. Besides ferrihydrite, soils from pretreatment and static anoxic conditions have features akin to Fe(II) bearing pyrite (FeS2; 8-14%). Whereas in the fluctuating and static oxic treatments, goethite, a crystalline Fe-oxide [α FeO(OH)] is the major Fe mineral. Comparison of the radial structure functions (RSFs) of soils under the four redox treatments with that of pure Fe minerals is presented in SI Figure S1B. A second shell feature indicates a similarity in the second coordination shell of Fe between 2.3 – 3.3 Å indicates similarity in Fe speciation between static anoxic and ferrihydrite whereas the fluctuating treatment and static oxic soils had increased crystallinity and paired more appropriately with goethite. This observation is in agreement with our wet chemistry data where the Fe concentrations for the fluctuating treatments closely resembled to that of the static oxic treatment.

**Table 1:** Bulk Fe K-edge EXAFS analysis. Linear combination fit (LCF) analysis results for the Fe K-edge EXAFS spectra of soils under the four redox treatments. Percentage of Fe species based on linear combination fitting of Fe EXAFS spectra using the following phases: ferrihydrite, goethite, pyrite, magnetite, siderite.

| Soil | ferrihydrite | goethite | pyrite | R-factor |
| --- | --- | --- | --- | --- |
| ----- % contribution ----- |  |  |  | (%) |
| pretreatment | **74.4<sup>‡</sup>** | 17.6 | 8.0 | 0.10 |
| Static anoxic | **86.7** | 0 | 13.3 | 0.31 |
| flux 4-day | **74.2** | 25.8 | 0 | 0.09 |
| flux 8-day | **73.1** | 26.9 | 0 | 0.15 |
| Static oxic | **58.5** | 41.5 | 0 | 0.20 |

<sup>‡</sup> Values in bold are majority components, as identified by linear combination fits

The decrease in AO-Fe concentrations during the fluctuating and static oxic treatments suggest a depletion of the kinetically labile Fe(III)-oxide pool in response to redox manipulation. On the other hand, the convergence of the DOC concentration around a similar value towards the end of the experiment for all the redox treatments indicates a potential equilibration point of the soils where the labile C sources possibly get exhausted and further points how the system is dependent on continued new C inputs. Prolonged anoxia in the soils further promotes the reductive dissolution of at least a fraction of SRO Fe oxides that might be present, thus releasing soluble Fe(II) which can then bind to functional groups of OM. We, therefore, observe a positive correlation between the reactive Fe-oxide phases and DOC under anoxic conditions indicating that Fe-oxides are instrumental in preserving OC via co-precipitation or direct chelation.^15^

The presence of crystalline long range ordered goethite (α-FeOOH) under fluctuating redox conditions can be explained by alternate dissolution and precipitation of the metastable ferrihydrite phase via thermodynamically favored Ostwald ripening.^54-56^ This is also in agreement to the results obtained from a redox oscillation study conducted on tropical soils of Hawaii.^6^ These alternate reductive Fe dissolution and precipitation mechanisms are often catalyzed by increased metabolic potential which could facilitate Fe-OM interactions. Elevated levels of soluble Fe(II) under anaerobic conditions potentially catalyze the reductive dissolution of crystalline Fe in presence of –O or –S containing organic ligands such as oxalate or thiol ^57-59^ that could, in turn, variably decompose or protect SOM via the formation of Fe-organic complexes or co-precipitates.^3, 60^ This is consistent with preferential interactions between Fe oxide minerals and reactive oxygen species (ROS) of aromatic moieties within polyfunctional SOM.^1, 2 61-63^ The observed increase in soluble Fe(II) concentrations during anoxic events could also be the result of bioreduction in presence of lignin-like compounds since we observe a shift in the LEF soil microbial community under reducing conditions (parallel study, data not shown). On the other hand, under oxic conditions, availability of Fe(III) oxides as a terminal electron acceptor^64^, increase in pH, or presence of ROS might have sustained CO_2_ production despite protective effects of Fe-C interactions.^1, 7^

### Short term redox shifts--effects on Fe and DOC chemistry

In addition to examining whole-experiment trends in 0.5 M HCl extractable Fe(II) and AO-Fe concentrations, we also assessed short-term effects (minutes to hours) of rapid redox changes within the fluctuating treatment soils (Figure 2). Since the trends for the flux 4-day and flux 8-day treatment soils were statistically similar, for simplicity, we present the average Fe(II) and AO-Fe concentrations of the two treatments combined in Figure 2. Soluble Fe(II) and AO-Fe concentrations increased considerably within 30 mins after a reduction event (switching from oxic to anoxic conditions). The opposite trend was observed during oxidation events, when the headspace was changed from anoxic to oxic, HCL and AO extractable Fe concentrations dropped dramatically. Similar to soluble Fe(II), water extractable DOC content also increased dramatically within 30 mins of switching from oxic to anoxic conditions (and vice versa) (Figure 2), implying that this DOC pool becomes available for respiration as early as 30 mins after change in redox condition. To our knowledge, these are some of the first evidence where we see such rapid changes in solution chemistry due to quick redox shifts. However, soils under these fluctuating conditions tend to exhibit dramatic chemical changes when exposed to opposite redox conditions even for short pulses as shown by our “quick” switch data. Specifically, for these short and fast redox events we observed that the frequency of low-redox events controls exchanges of C between mineral-sorbed and aqueous pools which in turn, affects whether the labile C is potentially lost as CO_2_ or via leaching. The rapid increase of labile Fe and DOC pools with the onset of anoxic conditions can be attributed to the reductive dissolution of crystalline Fe oxides leading to the accumulation of soluble Fe(II) and AO-Fe during the anoxic phases. The decrease in AO-Fe concentrations during the fluctuating and static oxic treatments suggest a depletion of the kinetically labile Fe(III)-oxide pool in response to redox manipulation. This correlation which is more dramatic during the short term redox switches is possibly due to either direct OM release from dissolving Fe(III) mineral phases or due to acidification since we observe a pH decline of ~ 0.2 units during the reduction event when DOC concentration increases almost three-fold (SI Figure S4).^7^

**Figure 2:**
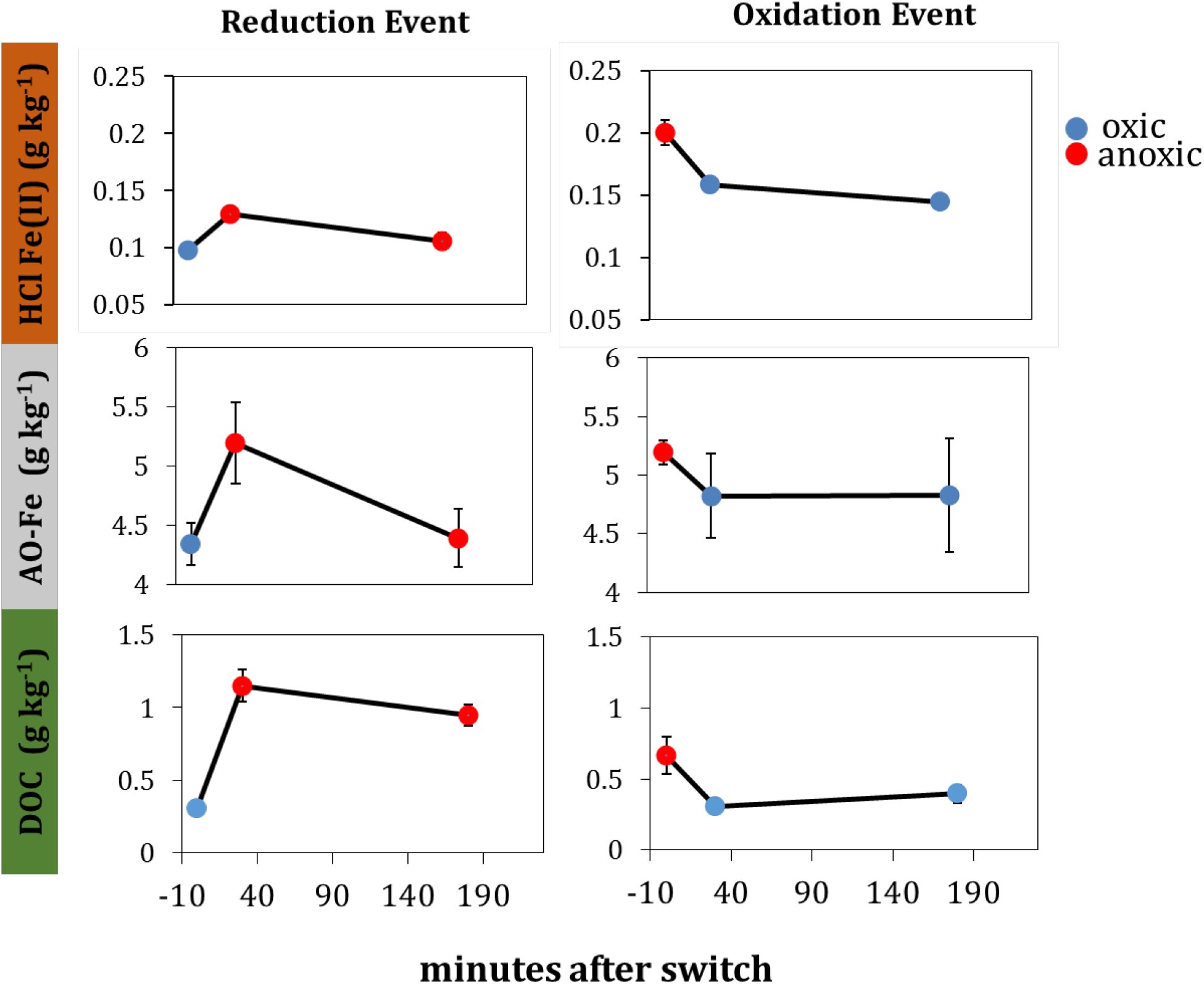
Changes in 0.5N HCl extractable Fe(II) [top], oxalate extractable Fe [middle] and WEDOC [bottom] concentrations (in g kg^−1^ soil) during rapid redox switches for reduction (oxic to anoxic) and oxidation (anoxic to oxic) events presented in left and right panels respectively. Blue and red circles indicate oxic and anoxic time points respectively. Error bars indicate standard error for three replicate measurements.

However, anaerobic microsites often instrumental in preferential protection of SOM are also susceptible to variable redox effects leading to a pulse DOC release (as shown by the shortterm switch data) and the remaining SOM being retained by metal-organic association primarily via complexation, sorption or co-precipitation in the soil as shown in previous studies.^65, 66^ Moreover, the rapid switch data suggests that redox shifts may promote higher nutrient and elemental cycling than static oxic (or anoxic) conditions via biotic or abiotic processes.

### Effects of redox shifts on the chemical composition of water extractable DOC

To test the dominant class of water extractable metabolites during redox changes, we investigated the chemical composition of the water extractable DOC pool via FT-ICR-MS technique.^67^ The FT-ICR-MS data show that although these soils began with the same metabolite composition (pretreatment), the fate of C (C speciation/composition) was dependent on redox conditions. Linear plots for nominal oxidation state of carbon (NOSC) for each of the redox regimes (Figures 3a-d) are created to aid in the identification of the relative distribution of the potential major organic C classes (i.e., lipids, proteins, lignin, carbohydrates, and condensed aromatics) under the different redox conditions.^68^ It is observed that the anoxic treatment is dominated by compounds with more reduced C values (more negative average NOSC, i.e. more reduced carbon) compared to the fluctuating and static oxic treatments (Figures 3a-d). A closer inspection of the slopes obtained from the trendlines of Figures 3a-d clearly show more negative values across static oxic (100% O2) to static anoxic (0% O2) treatments. Figure 3e summarizes our observation and illustrates how the slope for each treatment has a positive correlation with the percentage exposure to oxygen suggesting an increase in the abundance of oxidized compounds with increased O2 abundance. Under anaerobic conditions, microbial communities will leave behind energy poor compounds (low NOSC) as these compounds don’t provide the energy needed for microbial communities, as a result the bulk NOSC. This is referred to as thermodynamic preservation of carbon in anoxic environments, thereby shifting the average NOSC of the entire water-soluble C pool towards more negative values for soils under anaerobic conditions.^69^ Under oxic conditions, decomposition proceeds without thermodynamic limitations. Pairwise score plots between selected principal components (PCs) of flux 4-day and flux 8-day treatments show that the separation between the treatments was not significant for flux 4-day samples (Figure S2). The explained variance of each PC is shown in the corresponding diagonal cells (Figure S2). Hence, we subsequently created 2D score plots only for soils under flux 8-day treatment as a function of headspace composition (oxic vs anoxic). This created a significant difference for some samples depending on whether the sampling headspace was oxic or anoxic (Figure 4a). Interestingly, the most significant differences between oxic and anoxic soil compounds were observed early on in the experiment, suggesting that most active sites have been oxidized during that time in the experiment and with time, no significant effect was observed even upon switching redox conditions. To further understand the differences between oxic and anoxic treatments for flux-8 day, the VIP score was used as a measure of a variable’s importance in the PLS-DA model. It summarizes the contribution a variable makes to the model. The weights correspond to the percentage variation explained by the PLS-DA component in the model (Figure 4b). The number of terms in the sum depends on the number of PLS-DA components found to be significant in distinguishing the classes. The X axis indicates the VIP scores corresponding to each variable on the Y-axis. The compounds on the right indicate the factors with the highest VIP scores and thus are the most contributory variables in class discrimination in the PLS-DA model. We then plotted the O/C and H/C ratios of the most significant organic compounds (Figure 4c). Compounds that were significantly abundant in the anoxic samples had higher H/C and lower O/C values belonging to the lignin-like class of compounds (Figure 4c). On the other hand, compounds abundant in the oxic headspace mostly had high H/C and high O/C values and fit in the tannin- and cellulose-like region of the van-Krevelen diagram (Figure 4c). These results suggest that oxidation targets specific classes of compounds such as aromatic compounds and/or carbohydrates depending on the prevailing redox condition.

**Figure 3.**
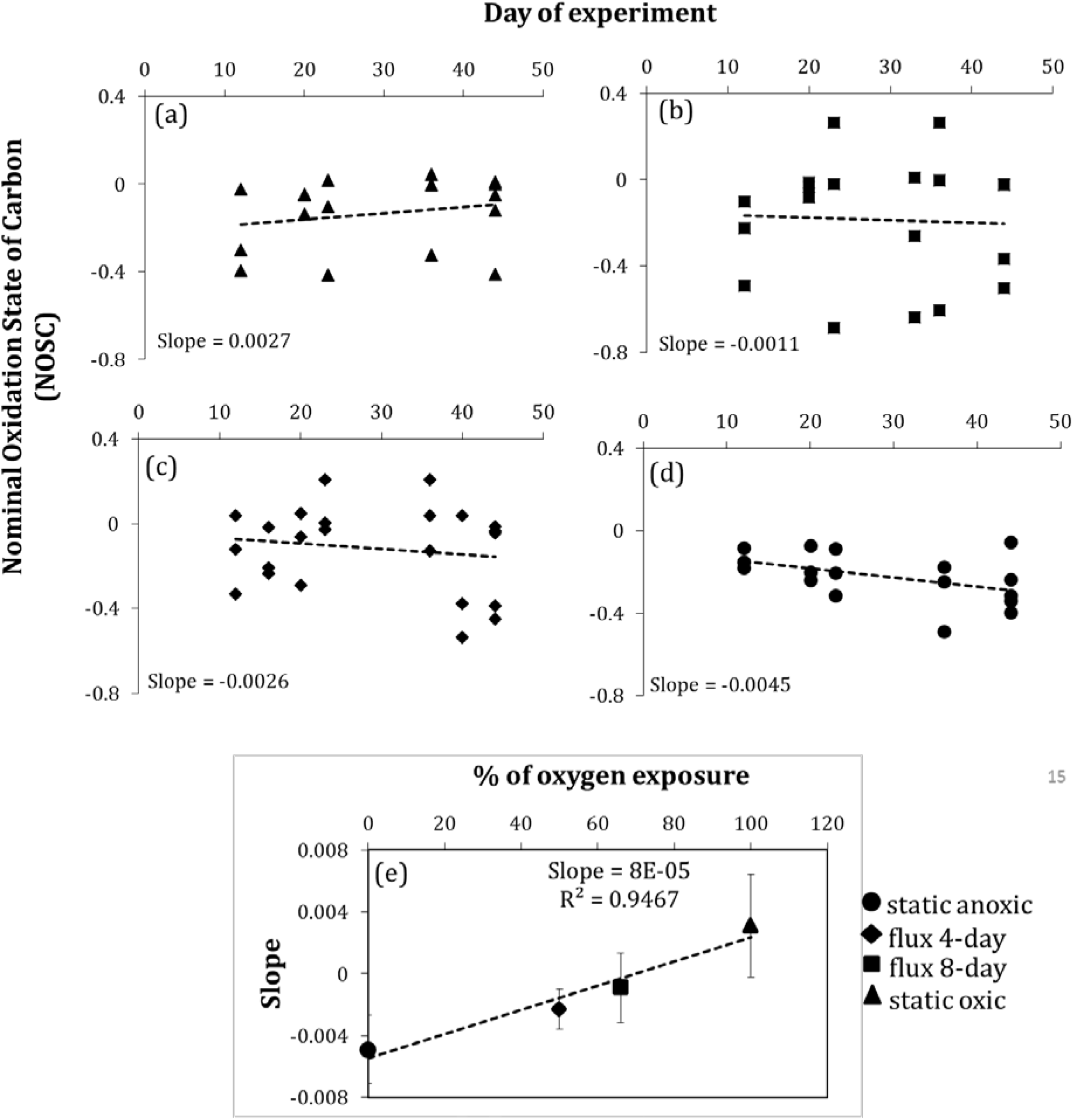
Fourier-transform ion-cyclotron mass spectrometry (FT-ICR-MS) analysis of water extractable metabolites. Average nominal oxidation state of carbon (NOSC) values of water soluble organic carbon fraction of a tropical forest soil from Puerto Rico over a 44-day redox incubation study. Plots a, b, c and d show the NOSC values over the 44-day for static oxic, flux 8-day, flux 4-day and static anoxic treatments respectively. Plot e shows how the slope of the lines (obtained from the trendlines in plots a-d) vary as the percent of oxygen changes across the four redox treatments.

**Figure 4.**
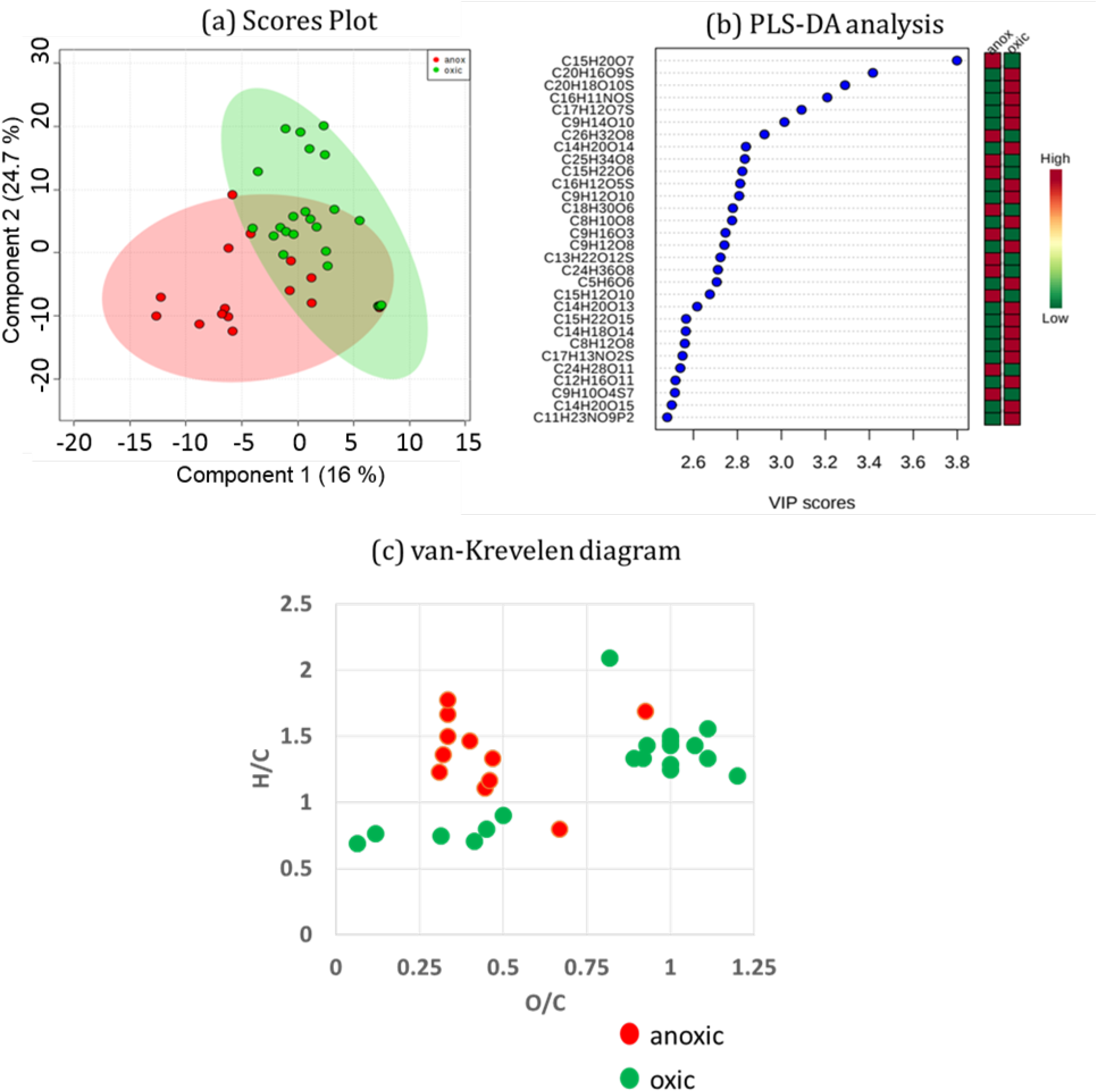
(a) 2D Score plot (component 1, 2) of anoxic (red circle) vs oxic (green circle) headspace in flux 8-day treatment. The dashed circle in the plot indicate later data points during the experiment; (b) important features identified by principal least square – discriminant analysis (PLS-DA). The colored boxes on the right indicate the relative concentrations of the corresponding metabolite in each group under anoxic or oxic conditions; and (c) van-Krevelen diagram (ratio of elemental H/C and O/C) of unique water extractable metabolites under oxic (green circle) and anoxic (red circle) conditions.

### Trace gas (CO_2_) emission under variable redox conditions

Geochemical changes measured by wet chemistry and molecular scale characterization studies (EXAFS and FT-ICR-MS) were complemented by macro scale measurements of the CO_2_ flux. ANOVA results showed that redox treatment significantly affected cumulative CO_2_ emission (P = 0.013), with cumulative CO_2_ flux highest in the static oxic treatment. Tukey’s tests suggest that the only significant pair-wise comparison was that cumulative CO_2_ flux was higher in the static oxic treatment than in the flux 8-day treatment (P = 0.009). Static oxic treatment also tended to increase cumulative CO_2_ flux relative to the static anoxic treatment (P =0.09) and the ‘flux 4-day” treatment (P = 0.11) according to Tukey’s tests. Concentrations measured during oxic time points during the fluctuating treatment incubations (days 12, 20, 36, and 44 for flux 4-day; and days 20, 33 and 44 for flux 8-day) behaved similarly to that of the static oxic treatment and had higher CO_2_ fluxes than anoxic sampling points (SI Figure S3a). However, the fluctuating treatments did not show a clear trend (SI Figure S3b). This is in agreement with previous studies which show CO_2_ production rates are often similar regardless of whether bulk soil Eh is low or high in LEF soils.^12, 34^

### Ecosystem Implications

Humid tropical soils of Puerto Rico are frequently subjected to alternating oxic and anoxic periods. This study has been built on the premise that predicted changes in global precipitation and temperature will cause a shift in the frequency, duration and degree of reduced conditions in these tropical soils, resulting in significantly altered mineral-organic interactions, thereby influencing tropical soil C cycling. Our study clearly shows how Fe-C coupling primarily controls redox-driven biogeochemistry in these soils that includes electron transfer reactions (i.e. oxidation of carbon and reduction of iron or other acceptors). The conceptual understanding gained by this research helps to identify key biogeochemical pathways, tipping points and parameters that are highly correlated with C retention or loss rates. Since the frequency of O2 depletion during a soil redox oscillation also shapes microbial community structure, functional stability and process rates, multiple C degradation pathways possibly co-occur in relatively short timeframes (similar to the fast redox switches), leading to a more rapid degradation of complex C substrates. Correspondingly, since C efflux from these soils is largely in the form of greenhouse gas, CO_2_, any alteration in the soil C emissions in response to changing redox can serve as a significant potential feedback to climate change. Therefore, to more accurately predict the ramifications of a warmer, drier, more stochastic neotropical biome, a mechanistic understanding of how different redox oscillation patterns affect biogeochemical mediation of C retention and loss pathways is critical and this data needs to eventually be reflected in numerical models.^70^ Our findings, therefore, highlight the importance of why redox cycling needs to be incorporated as a non-linear predictive variable into the emerging image of soil C stabilization as an ecosystem property.^71^ The mechanistic understanding of Fe-OM interactions obtained from this study will directly benefit improvements to predictive mathematical models that forecast future tropical soil carbon balance in future ecosystem scale studies.

## ASSOCIATED CONTENT

### SUPPORTING INFORMATION

The Supporting Information is available free of charge on the ACS Publications website. Contains details on experimental methods and four figures.

#### Notes

The authors declare no competing financial interest.

## ACKNOWLEDGEMENTS

We thank Daniel Nilson, Elizabeth Green, Jessica Wollard, Shalini Mabery, Rachel Neurath, Keith Morrison, Christopher Ward, Jeffery Kimbrel, Mona Hwang and Feliza Bourguet for assistance in the laboratory. This project was supported by a US Department of Energy Early Career Award to J. Pett-Ridge (SCW1478) administered by the Office of Biological and Environmental Research, Genomic Sciences Program. Work at LLNL was performed under the auspices of the U.S. Department of Energy under Contract DE-AC52-07NA27344. Synchrotron work was performed at beamlines 5.3.2.2 and 11.2.2 of the Advanced Light Source, which is a DOE Office of Science User Facility under contract no. DE-AC02-05CH11231. FTICRMS was performed using EMSL, a DOE Office of Science User Facility sponsored by BER and located at Pacific Northwest National Laboratory (PNNL).

## Supporting Information

Contain details on linear combination fit analyses of Fe K-edge EXAFS data and four figures. This material is available free of charge via the Internet at http://pubs.acs.org.

## Methods

<u>Linear Combination Fitting:</u> Combinatorial linear combination fitting analyses were performed using Athena software on k^3^-weighted Fe EXAFS spectra from 2-12. Sums were floated and all components were limited between 0 and 1. Number of standards was limited to 8 and no minimum contribution of standard was required. Representative fits of three samples are shown in Figure S6. Many of the top combinatorial fits were statistically indistinguishable by the Hamilton test^1^, so the top 10 fits for each sample were averaged, and the reported R factor represents the goodness of fit. The top 10 combinatorial fits and their average are shown in Figure S1.

## Supplementary Figures

**Figure S1.**
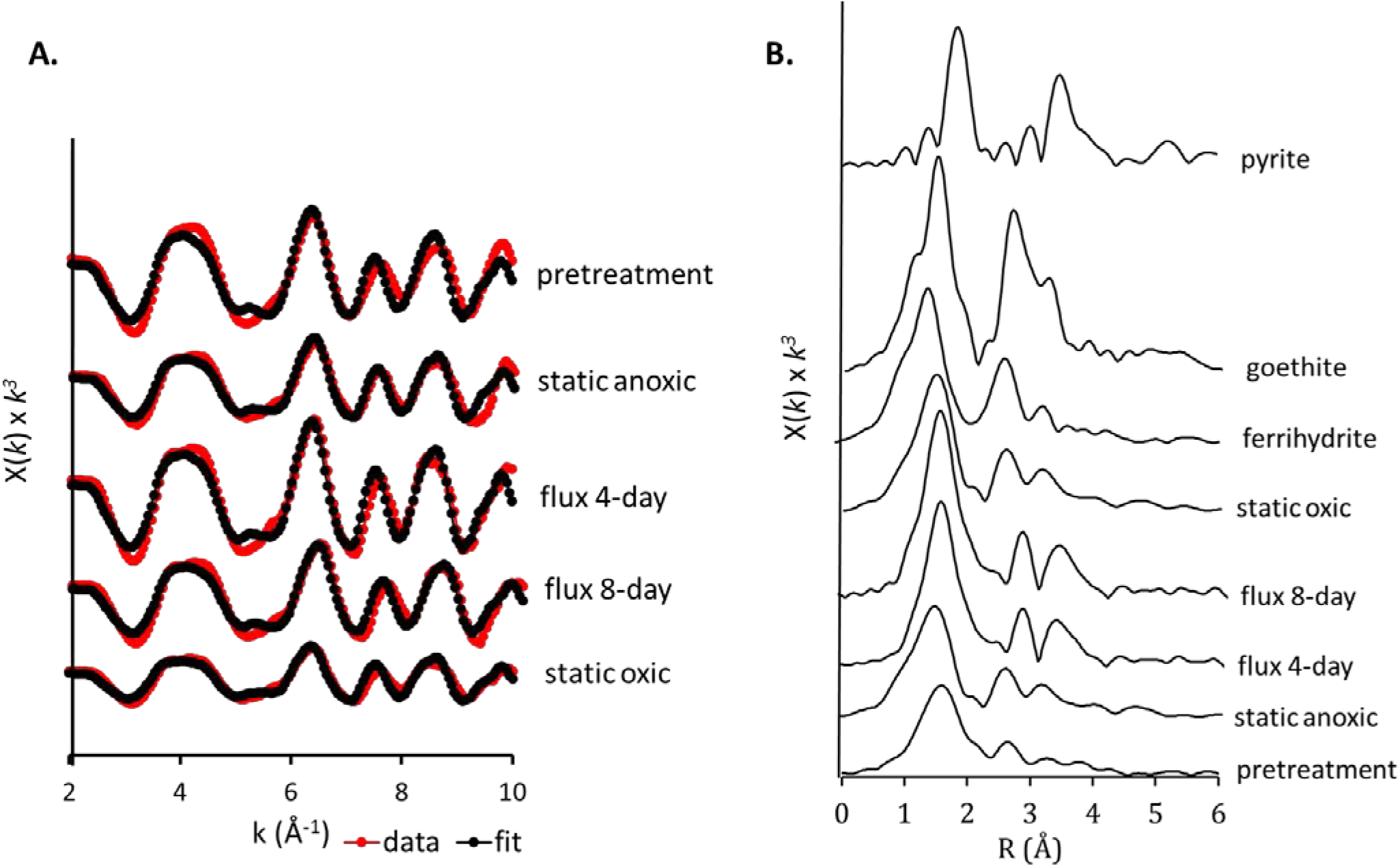
(A) EXAFS analysis. *k^3^*-weighted EXAFS spectra and fits of soils from Puerto Rico under pretreatment and four redox conditions (top to bottom). Solid (red) and dotted (black) lines indicate the experimental and fitted Fe K-edge EXAFS spectra, respectively. (B) Radial structure functions (RSF) of soils under the different redox treatments and model Fe minerals commonly found in Puerto Rican soils are reported. Pretreatment and static anoxic soils show more similarities to that of amorphous ferrihydrite and pyrite whereas soils under static oxic and fluctuating redox conditions are more similar to crystalline goethite.

**Figure S2.**
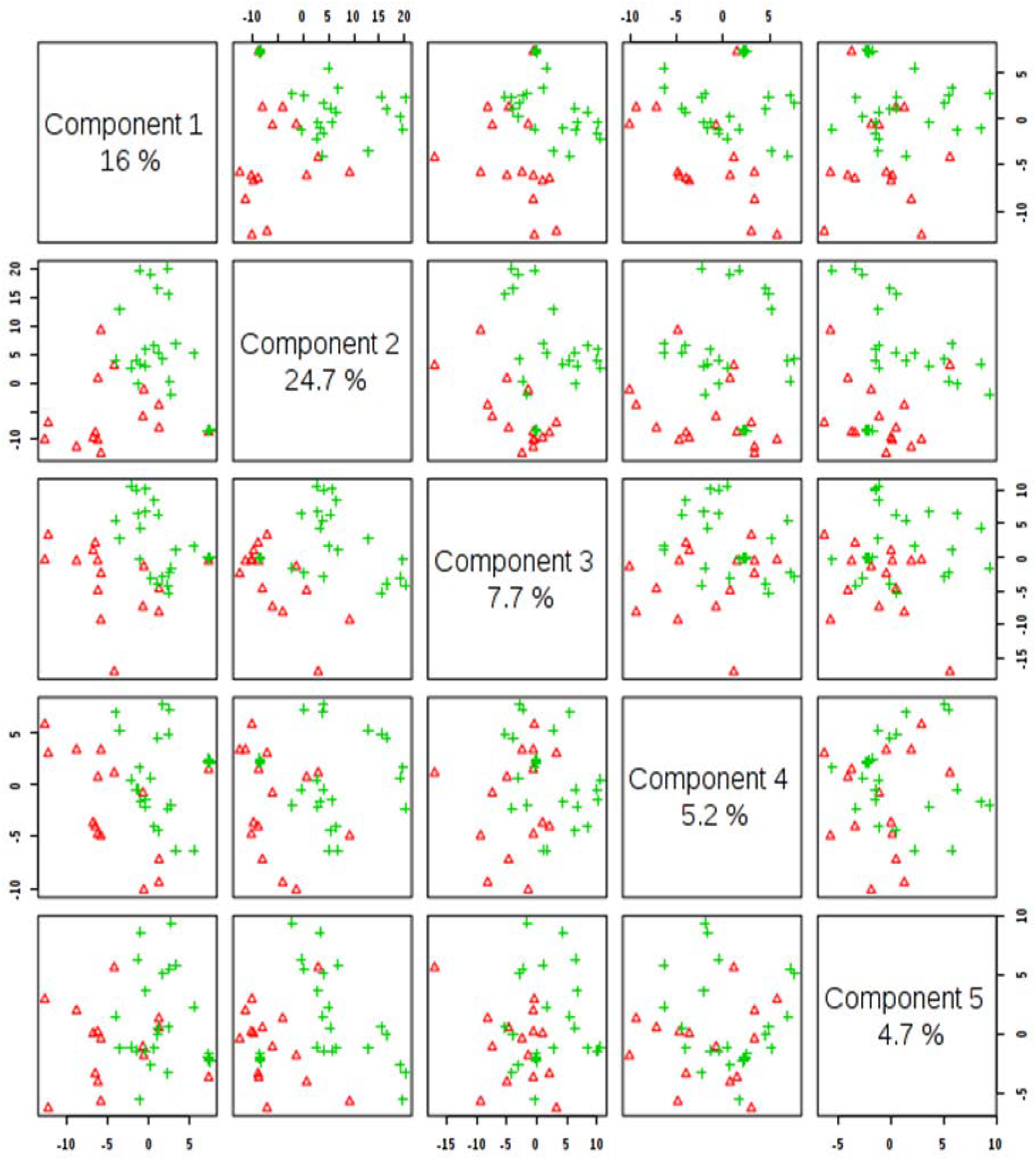
Pairwise score plots between the selected principal components (PCs) of flux 4-day (shown in red) and flux8-day (shown in green) treatments in tropical soils of Puerto Rico. The explained variance of each PC is shown in the corresponding diagonal cell. Separation between treatments was not significant for flux 4-day samples.

**Figure S3.**
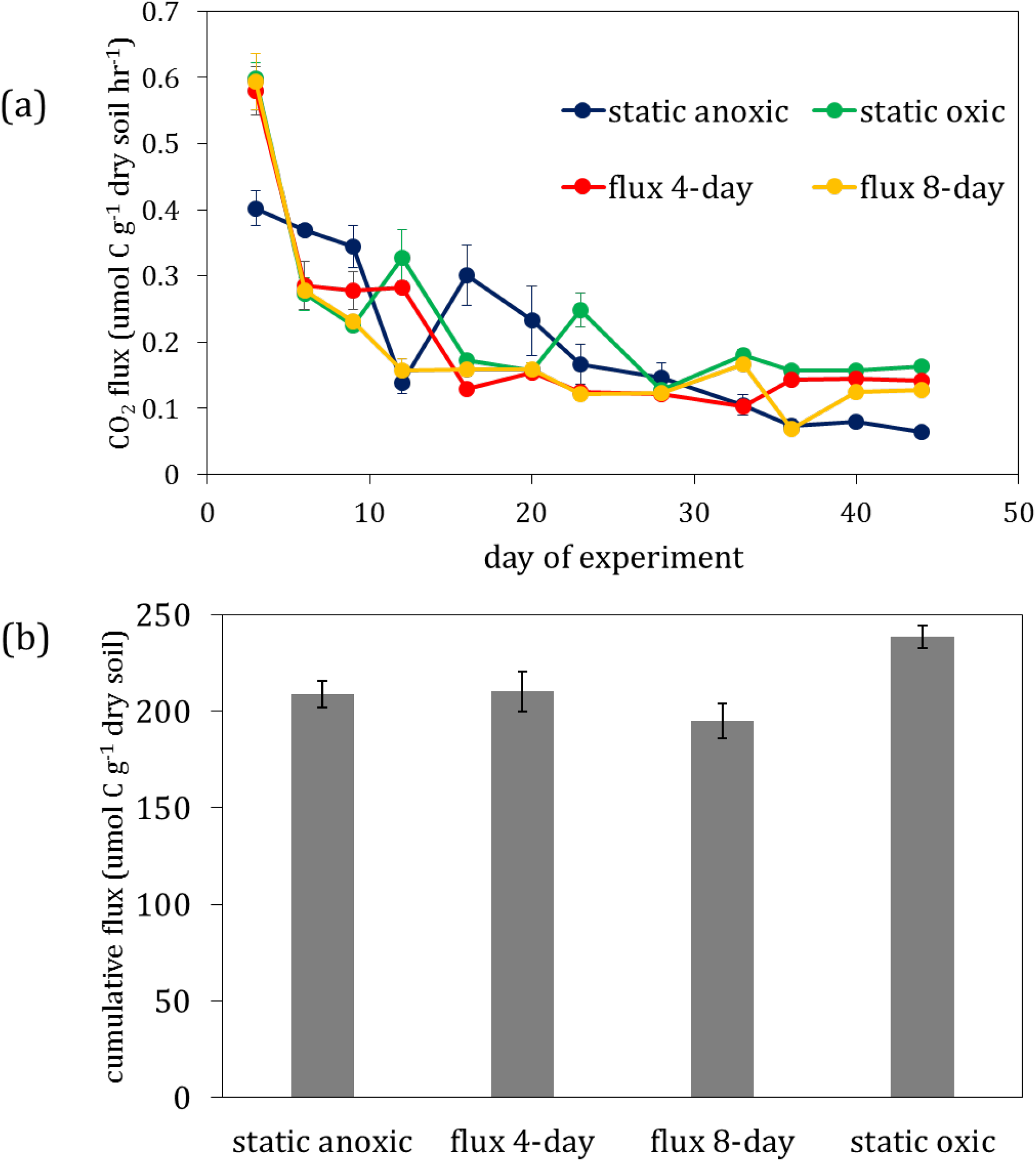
(a) Instantaneous CO_2_ production and (b) cumulative CO_2_ production over the 44-day incubation on a tropical forest soil from Puerto Rico under four redox treatments static oxic (green), flux 8-day (yellow), flux 4-day (red), static anoxic (blue). Error bars indicate standard error for five replicate measurements.

**Figure S4.**
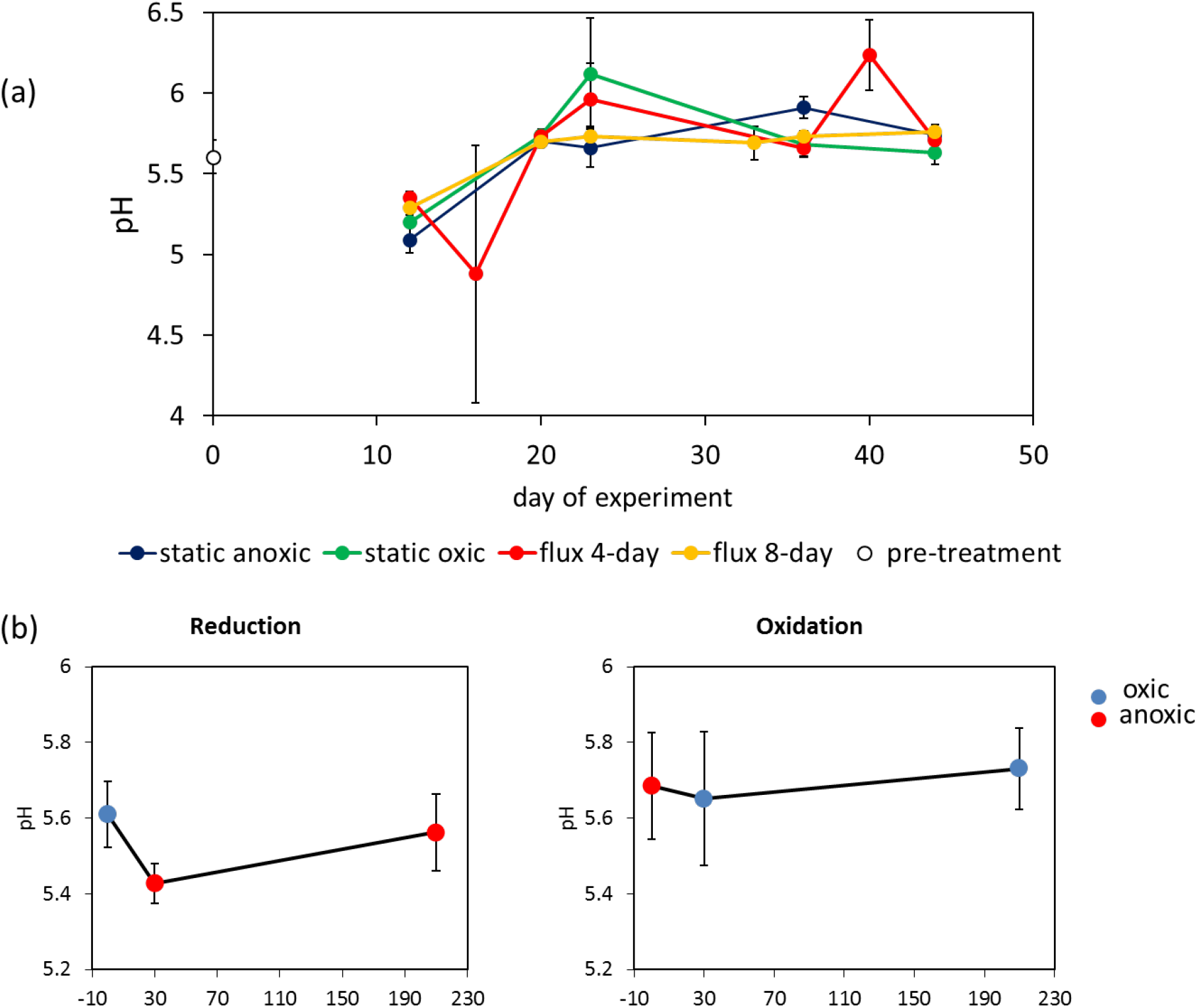
(a) pH changes over the 44-day incubation on a tropical forest soil from Puerto Rico under four redox treatments. pH values for static anoxic, static oxic, flux 4-day, flux 8-day treatments are shown in blue, green, red and yellow respectively compared to that of the initial pretreatment pH (black circle). Error bars indicate standard error for three replicate measurements. (b) pH changes during rapid redox switches for reduction (oxic to anoxic) and oxidation (anoxic to oxic) events presented in left and right panels respectively. Blue and red circles indicate oxic and anoxic time points respectively. Error bars indicate standard error for three replicate measurements.

## REFERENCES

1. Hall, S. J.; Silver, W. L., Iron oxidation stimulates organic matter decomposition in humid tropical forest soils. Global change biology 2013, 19, (9), 2804–2813.

2. Hall, S. J.; Silver, W. L.; Timokhin, V. I.; Hammel, K. E., Lignin decomposition is sustained under fluctuating redox conditions in humid tropical forest soils. Global change biology 2015, 21, (7), 2818–2828.

3. Hall, S. J.; Silver, W. L.; Timokhin, V. I.; Hammel, K. E., Iron addition to soil specifically stabilized lignin. Soil Biology and Biochemistry 2016, 98, 95–98.

4. Liptzin, D.; Silver, W.; Detto, M., Temporal Dynamics in Soil Oxygen and Greenhouse Gases in Two Humid Tropical Forests. Ecosystems 2011, 14, (2), 171–182.

5. Silver, W. L.; Lugo, A. E.; Keller, M., Soil oxygen availability and biogeochemistry along rainfall and topographic gradients in upland wet tropical forest soils. Biogeochemistry 1999, 44, (3), 301–328.

6. Thompson, A.; Chadwick, O. A.; Rancourt, D. G.; Chorover, J., Iron-oxide crystallinity increases during soil redox oscillations. Geochimica et Cosmochimica Acta 2006, 70, (7), 1710–1727.

7. Thompson, A.; Chadwick, O. A.; Boman, S.; Chorover, J., Colloid mobilization during soil iron redox oscillations. Environmental Science & Technology 2006, 40, (18), 5743–5749.

8. Thompson, A.; Rancourt, D. G.; Chadwick, O. A.; Chorover, J., Iron solid-phase differentiation along a redox gradient in basaltic soils. Geochimica et Cosmochimica Acta 2011, 75, (1), 119–133.

9. Tishchenko, V.; Meile, C.; Scherer, M. M.; Pasakarnis, T. S.; Thompson, A., Fe2+ catalyzed iron atom exchange and re-crystallization in a tropical soil. Geochimica et Cosmochimica Acta 2015, 148, (0), 191–202.

10. Dubinsky, E. A.; Silver, W. L.; Firestone, M. K., Tropical forest soil microbial communities couple iron and carbon biogeochemistry. Ecology 2010, 91, (9), 2604–2612.

11. Ponnamperuma, F.; Tianco, E. M.; Loy, T., Redox equilibria in flooded soils: I. The iron hydroxide systems. Soil Science 1967, 103, (6), 374–382.

12. Pett-Ridge, J. Rapidly fluctuating redox regimes frame the ecology of microbial communities and their biogeochemical function in a humid tropical soil. University of California, Berkeley, Berkeley, 2005.

13. Reddy, K.; Patrick Jr, W., Effect of alternate aerobic and anaerobic conditions on redox potential, organic matter decomposition and nitrogen loss in a flooded soil. Soil Biology and Biochemistry 1975, 7, (2), 87–94.

14. Ginn, B. R.; Meile, C.; Wilmoth, J.; Tang, Y.; Thompson, A., Rapid iron reduction rates are stimulated by high-amplitude redox fluctuations in a tropical forest soil. 2017.

15. Lalonde, K.; Mucci, A.; Ouellet, A.; Gelinas, Y., Preservation of organic matter in sediments promoted by iron. Nature 2012, 483, (7388), 198–200.

16. Parr, J.; Reuszer, H., Organic matter decomposition as influenced by oxygen level and method of application to soil. Soil Science Society of America Journal 1959, 23, (3), 214–216.

17. Kaiser, K.; Guggenberger, G., Sorptive stabilization of organic matter by microporous goethite: sorption into small pores vs. surface complexation. European Journal of Soil Science 2007, 58, (1), 45–59.

18. Kaiser, K.; Mikutta, R.; Guggenberger, G., Increased Stability of Organic Matter Sorbed to Ferrihydrite and Goethite on Aging. Soil Sci Soc Am J 2007, 71, (3), 711–719.

19. Koegel-Knabner, I.; Guggenberger, G.; Kleber, M.; Kandeler, E.; Kalbitz, K.; Scheu, S.; Eusterhues, K.; Leinweber, P., Organo-mineral associations in temperate soils: Integrating biology, mineralogy, and organic matter chemistry. Journal of Plant Nutrition and Soil Science-Zeitschrift Fur Pflanzenernahrung Und Bodenkunde 2008, 171, (1), 61–82.

20. Lutzow, M. v.; Kogel-Knabner, I.; Ekschmitt, K.; Matzner, E.; Guggenberger, G.; Marschner, B.; Flessa, H., Stabilization of organic matter in temperate soils: mechanisms and their relevance under different soil conditions - a review. European Journal of Soil Science 2006, 57, (4), 426–445.

21. Wagai, R.; Mayer, L. M., Sorptive stabilization of organic matter in soils by hydrous iron oxides. Geochimica et Cosmochimica Acta 2007, 71, (1), 25–35.

22. Amstaetter, K.; Borch, T.; Kappler, A., Influence of humic acid imposed changes of ferrihydrite aggregation on microbial Fe(III) reduction. Geochimica et Cosmochimica Acta 2012, 85, (0), 326–341.

23. Cismasu, A. C.; Michel, F. M.; Tcaciuc, A. P.; Tyliszczak, T.; Brown, G. E., Composition and structural aspects of naturally occurring ferrihydrite. C. R. Geosci. 2011, 343, (2–3), 210–218.

24. Pedrot, M.; Le Boudec, A.; Davranche, M.; Dia, A.; Henin, O., How does organic matter constrain the nature, size and availability of Fe nanoparticles for biological reduction? J Colloid Interf Sci 2011, 359, (1), 75–85.

25. Mikutta, R.; Kleber, M.; Torn, M. S.; Jahn, R., Stabilization of soil organic matter: Association with minerals or chemical recalcitrance? Biogeochemistry 2006, 77, (1), 25–56.

26. Torn, M. S.; Trumbore, S. E.; Chadwick, O. A.; Vitousek, P. M.; Hendricks, D. M., Mineral control of soil organic carbon storage and turnover. Nature 1997, 389, (6647), 170–173.

27. Liptzin, D.; Silver, W. L., Effects of carbon additions on iron reduction and phosphorus availability in a humid tropical forest soil. Soil Biology and Biochemistry 2009, 41, (8), 1696–1702.

28. Chacon, N.; Silver, W. L.; Dubinsky, E. A.; Cusack, D. F., Iron Reduction and Soil Phosphorus Solubilization in Humid Tropical Forests Soils: The Roles of Labile Carbon Pools and an Electron Shuttle Compound. Biogeochemistry 2006, 78, (1), 67–84.

29. Buettner, S. W.; Kramer, M. G.; Chadwick, O. A.; Thompson, A., Mobilization of colloidal carbon during iron reduction in basaltic soils. Geoderma 2014, 221, 139–145.

30. Change, I. C., The Fourth Assessment Report of the Intergovernmental Panel on Climate Change. Geneva, Switzerland 2007.

31. Bouskill, N. J.; Lim, H. C.; Borglin, S.; Salve, R.; Wood, T. E.; Silver, W. L.; Brodie, E. L., Pre-exposure to drought increases the resistance of tropical forest soil bacterial communities to extended drought. The ISME journal 2013, 7, (2), 384.

32. Bouskill, N. J.; Wood, T. E.; Baran, R.; Hao, Z.; Ye, Z.; Bowen, B. P.; Lim, H. C.; Nico, P. S.; Holman, H.-Y.; Gilbert, B., Belowground response to drought in a tropical forest soil. II. Change in microbial function impacts carbon composition. Frontiers in microbiology 2016, 7, 323.

33. Bouskill, N. J.; Wood, T. E.; Baran, R.; Ye, Z.; Bowen, B. P.; Lim, H.; Zhou, J.; Nostrand, J. D. V.; Nico, P.; Northen, T. R., Belowground response to drought in a tropical forest soil. I. Changes in microbial functional potential and metabolism. Frontiers in microbiology 2016, 7, 525.

34. DeAngelis, K. M.; Silver, W. L.; Thompson, A. W.; Firestone, M. K., Microbial communities acclimate to recurring changes in soil redox potential status. Environmental Microbiology 2010, 12, (12), 3137–3149.

35. Pett-Ridge, J.; Petersen, D. G.; Nuccio, E.; Firestone, M. K., Influence of oxic/anoxic fluctuations on ammonia oxidizers and nitrification potential in a wet tropical soil. FEMS Microbiology Ecology 2013, 85, (1), 179–194.

36. Pett-Ridge, J.; Silver, W. L.; Firestone, M. K., Redox Fluctuations Frame Microbial Community Impacts on N-cycling Rates in a Humid Tropical Forest Soil. Biogeochemistry 2006, V81, (1), 95–110.

37. Teh, Y. A.; Silver, W. L.; Conrad, M. E., Oxygen effects on methane production and oxidation in humid tropical forest soils. Global Change Biology 2005, 11, (8), 1283–1297.

38. Templer, P. H.; Silver, W. L.; Pett-Ridge, J.; DeAngelis, K.; Firestone, M. K., Plant and microbial controls on nitrogen retention and loss in a humid tropical forest. Ecology 2008, 89, (11), 3030–3040.

39. Wood, T. E.; Silver, W. L., Strong spatial variability in trace gasdynamics following experimental drought in a humid tropical forest. Global Biogeochemical Cycles 2012, 26, (3).

40. Pett-Ridge, J.; Firestone, M. K., Redox fluctuation structures microbial communities in a wet tropical soil. Appl. Environ. Microbiol. 2005, 71, (11), 6998–7007.

41. White, A. F.; Blum, A. E.; Schulz, M. S.; Vivit, D. V.; Stonestrom, D. A.; Larsen, M.; Murphy, S. F.; Eberl, D., Chemical weathering in a tropical watershed, Luquillo Mountains, Puerto Rico: I. Long-term versus short-term weathering fluxes. Geochimica et Cosmochimica Acta 1998, 62, (2), 209–226.

42. McKeague, J.; Day, J., Dithionite-and oxalate-extractable Fe and Al as aids in differentiating various classes of soils. Canadian journal of soil science 1966, 46, (1), 13–22.

43. Stookey, L. L., Ferrozine---a new spectrophotometric reagent for iron. Analytical chemistry 1970, 42, (7), 779–781.

44. Webb, S., SIXpack: a graphical user interface for XAS analysis using IFEFFIT. Physica scripta 2005, 2005, (T115), 1011.

45. Ravel, B.; Newville, M., ATHENA, ARTEMIS, HEPHAESTUS: data analysis for X-ray absorption spectroscopy using IFEFFIT. Journal of synchrotron radiation 2005, 12, (4), 537–541.

46. Tfaily, M. M.; Chu, R. K.; Toyoda, J.; Tolić, N.; Robinson, E. W.; Paša-Tolić, L.; Hess, N. J., Sequential extraction protocol for organic matter from soils and sediments using high resolution mass spectrometry. Analytica chimica acta 2017, 972, 54–61.

47. Tfaily, M. M.; Hamdan, R.; Corbett, J. E.; Chanton, J. P.; Glaser, P. H.; Cooper, W. T., Investigating dissolved organic matter decomposition in northern peatlands using complimentary analytical techniques. Geochimica et Cosmochimica Acta 2013, 112, (0), 116–129.

48. Tfaily, M. M.; Hodgkins, S.; Podgorski, D. C.; Chanton, J. P.; Cooper, W. T., Comparison of dialysis and solid-phase extraction for isolation and concentration of dissolved organic matter prior to Fourier transform ion cyclotron resonance mass spectrometry. Analytical and bioanalytical chemistry 2012, 404, (2), 447–457.

49. Tfaily, M. M.; Podgorski, D. C.; Corbett, J. E.; Chanton, J. P.; Cooper, W. T., Influence of acidification on the optical properties and molecular composition of dissolved organic matter. Analytica chimica acta 2011, 706, (2), 261–267.

50. Tolić, N.; Liu, Y.; Liyu, A.; Shen, Y.; Tfaily, M. M.; Kujawinski, E. B.; Longnecker, K.; Kuo, L.-J.; Robinson, E. W.; Paša-Tolić, L., Formularity: Software for Automated Formula Assignment of Natural and Other Organic Matter from Ultrahigh-Resolution Mass Spectra. Analytical Chemistry 2017, 89, (23), 12659–12665.

51. LaRowe, D. E.; Van Cappellen, P., Degradation of natural organic matter: a thermodynamic analysis. Geochimica et Cosmochimica Acta 2011, 75, (8), 2030–2042.

52. Xia, J.; Sinelnikov, I. V.; Han, B.; Wishart, D. S., MetaboAnalyst 3.0—making metabolomics more meaningful. Nucleic acids research 2015, 43, (W1), W251–W257.

53. Fredrickson, J. K.; Zachara, J. M.; Kennedy, D. W.; Dong, H.; Onstott, T. C.; Hinman, N. W.; Li, S.-m., Biogenic iron mineralization accompanying the dissimilatory reduction of hydrous ferric oxide by a groundwater bacterium. Geochimica et Cosmochimica Acta 1998, 62, (19), 3239–3257.

54. Deelman, J., Breaking Ostwald’s rule. Chemie Der Erde-Geochemistry 2001, 61, (3), 224–235.

55. Hansel, C. M.; Benner, S. G.; Neiss, J.; Dohnalkova, A.; Kukkadapu, R. K.; Fendorf, S., Secondary mineralization pathways induced by dissimilatory iron reduction of ferrihydrite under advective flow. Geochimica et Cosmochimica Acta 2003, 67, (16), 2977–2992.

56. Williams, A. G.; Scherer, M. M., Spectroscopic evidence for Fe (II)−Fe (III) electron transfer at the iron oxide−water interface. Environmental science & technology 2004, 38, (18), 4782–4790.

57. Kostka, J. E.; Luther, G. W., Partitioning and speciation of solid phase iron in saltmarsh sediments. Geochimica et Cosmochimica Acta 1994, 58, 1701–1710.

58. Sulzberger, B.; Suter, D.; Siffert, C.; Banwart, S.; Stumm, W., Dissolution of Fe(III)(hydr)oxides in natural waters; laboratory assessment on the kinetics controlled by surface coordination. Marine Chemistry 1989, 28, (1–3), 127–144.

59. Bhattacharyya, A.; Stavitski, E.; Dvorak, J.; Martínez, C. E., Redox interactions between Fe and cysteine: Spectroscopic studies and multiplet calculations. Geochimica et Cosmochimica Acta 2013, 122, 89–100.

60. Roden, E. E., Geochemical and microbiological controls on dissimilatory iron reduction. C. R. Geosci. 2006, 338, (6), 456–467.

61. Gu, B.; Schmitt, J.; Chen, Z.; Liang, L.; McCarthy, J. F., Adsorption and desorption of different organic matter fractions on iron oxide. Geochimica et Cosmochimica Acta 1995, 59, (2), 219–229.

62. Kaiser, K., Sorption of natural organic matter fractions to goethite (α-FeOOH): effect of chemical composition as revealed by liquid-state 13 C NMR and wet-chemical analysis. Organic Geochemistry 2003, 34, (11), 1569–1579.

63. Riedel, T.; Zak, D.; Biester, H.; Dittmar, T., Iron traps terrestrially derived dissolved organic matter at redox interfaces. Proceedings of the National Academy of Sciences 2013, 110, (25), 10101–10105.

64. Sutton-Grier, A. E.; Keller, J. K.; Koch, R.; Gilmour, C.; Megonigal, J. P., Electron donors and acceptors influence anaerobic soil organic matter mineralization in tidal marshes. Soil Biology and Biochemistry 2011, 43, (7), 1576–1583.

65. Keiluweit, M.; Wanzek, T.; Kleber, M.; Nico, P.; Fendorf, S., Anaerobic microsites have an unaccounted role in soil carbon stabilization. Nature communications 2017, 8, (1), 1771.

66. Bhattacharyya, A.; Schmidt, M. P.; Stavitski, E.; Martínez, C. E., Iron speciation in peats: Chemical and spectroscopic evidence for the co-occurrence of ferric and ferrous iron in organic complexes and mineral precipitates. Organic Geochemistry 2018, 115, 124–137.

67. Tfaily, M. M.; Chu, R. K.; Tolić, N.; Roscioli, K. M.; Anderton, C. R.; Paša-Tolić, L.; Robinson, E. W.; Hess, N. J., Advanced solvent based methods for molecular characterization of soil organic matter by high-resolution mass spectrometry. Analytical chemistry 2015, 87, (10), 5206–5215.

68. Kim, S.; Kramer, R. W.; Hatcher, P. G., Graphical method for analysis of ultrahigh-resolution broadband mass spectra of natural organic matter, the van Krevelen diagram. Analytical Chemistry 2003, 75, (20), 5336–5344.

69. Boye, K.; Noël, V.; Tfaily, M. M.; Bone, S. E.; Williams, K. H.; Bargar, J. R.; Fendorf, S., Thermodynamically controlled preservation of organic carbon in floodplains. Nature Geoscience 2017, 10, (6).

70. Treseder, K. K.; Balser, T. C.; Bradford, M. A.; Brodie, E. L.; Dubinsky, E. A.; Eviner, V. T.; Hofmockel, K. S.; Lennon, J. T.; Levine, U. Y.; MacGregor, B. J., Integrating microbial ecology into ecosystem models: challenges and priorities. Biogeochemistry 2012, 109, (1–3), 7–18.

71. Keiluweit, M.; Bougoure, J. J.; Nico, P. S.; Pett-Ridge, J.; Weber, P. K.; Kleber, M., Mineral protection of soil carbon counteracted by root exudates. Nature Climate Change 2015, 5, (6), 588–595.

## Reference

(1) Downward, L.; Booth, C. H.; Lukens, W. W.; Bridges, F. A variation of the F-test for determining statistical relevance of particular parameters in EXAFS fits. AIP Conf. Proc. 2007, 882, 129–131.

